# An algorithm for template-based prediction of secondary structures of individual RNA sequences

**DOI:** 10.1101/171108

**Authors:** Josef Pánek, Martin Černý

## Abstract

While understanding the structure of RNA molecules is vital for deciphering their functions, determining RNA structures experimentally is exceptionally hard. At the same time, extant approaches to computational RNA structure prediction have limited applicability and reliability. In this paper we provide a method to solve a simpler yet still biologically relevant problem: prediction of secondary RNA structure using structure of different molecules as a template.

Our method identifies conserved and unconserved subsequences within an RNA molecule. For conserved subsequences, the template structure is directly transferred into the generated structure and combined with de-novo predicted structure for the unconserved subsequences with low evolutionary conservation. The method also determines, when the generated structure is unreliable.

The method is validated using experimentally identified structures. The accuracy of the method exceeds that of classical prediction algorithms and constrained prediction methods. This is demonstrated by comparison using large number of heterogeneous RNAs. The presented method is fast and robust, and useful for various applications requiring knowledge of secondary structures of individual RNA sequences.

## INTRODUCTION

Experimental identification of RNA structures is extremely technically demanding. Therefore, computational predictions of RNA secondary structures are frequently employed as proxies for native structures. There is plenty of heterogeneous prediction methods, reviewed in Mathews et al. 2006 (1) for the free energy minimization and dynamic programing techniques, and in Gardner et al. 2004 (2) for comparative methods. However, known methods are unreliable namely for longer sequences (approx. > 150 nucleotides) and more complex structures, e.g. those that contain longer single-stranded segments. This is owing to the extreme theoretical complexity of the prediction.

Nevertheless, the number of experimentally identified RNA structures is growing in spite of the technical demands. These structures are available as potential templates to generate secondary structures of uncharacterized but related RNA sequences. In principle, template-based prediction can be treated as constrained prediction, which is supported by several methods, e.g. the RNA Vienna Package (3), RNAstructure (4) and Locarna (5). However, the conversion of the template into a structural constraint for Locarna and RNAstructure is not trivial, as the template and the query sequence frequently have different lengths. Locarna produces a consensus structure different from an individual structure that cannot be mapped directly. RNAstructure requires multiple input sequences as input otherwise it predicts either MFE structure or a set of probable structures. The only directly applicable method is thus via RNA Vienna Package.

It includes an utility, refold.pl, that can be used for conversion of secondary structures into an RNAfold constraint through a sequence alignment. The constraint is then used for RNAfold constrained prediction.

A different approach to the template-based prediction was adopted for ribosomal RNAs (6). The rRNA structure can not be predicted by available methods due to their lengths and complexity. The method uses known rRNA structures as templates for comparative prediction of homologous sequences but it requires extensive manual input and is slow.

We adopted a principally different approach: our method generates RNA secondary structure for the molecule under investigation directly from the template structure. The reliability of the generated structure is then evaluated by a bootstrapping scheme. We show that the proposed method achieves high-quality predictions for sequences where a structure for a putative homolog is known, including a number of sequences that are intractable by current prediction software.

An utility based on the method is available on request from the authors.

## MATERIALS AND METHODS

In this section, we first describe the proposed prediction method and then deal with the evaluation methodology.

### Generating a structure

Input to the template-based prediction task consists of a template sequence, the corresponding template structure and a query sequence. The task is to predict the query structure of the query sequence.

In terms of folding space, we have a subspace of the complete folding space of the query sequence. The subspace contains all possible structures of evolutionarily unconserved segments of the query structure, while the structure of evolutionarily conserved segments is taken from the template and kept fixed. The solution over this subspace is in principle easier than over the complete space, and can be found by determining the optimal structure of the unconserved segments. It is accomplished as follows:

I. A pairwise alignment of the query and template sequences is computed with ClustalW2 (with default parameters except for GAPOPEN = 7 a GAPEXT = 0.5) (7). The alignment can be treated as two functions: *A_q_* maps positions in the query sequence to positions in the template sequence and *A_t_* maps from template to query sequence. An example alignment is shown in Table S1.
II. The template structure is mapped onto the query sequence, producing *intermediate structure*. The intermediate structure preserves base pairs that the alignment maps to complementary nucleotides and marks all other bases as unpaired. More precisely, for each position *p* in the query sequence, there are four possibilities:

1. *A_q_(p)* is a gap,
2. *A_q_(p)* is not paired in the template structure,
3. *A_q_(p)* is paired to position r, but *A_t_(r)* is either a gap or a non-canonical pair for *p*,
4. *A_q_(p)* is paired to position *r* and *A_t_(r)* is a canonical pair for *p* In cases 1-3 the intermediate query structure marks *p* as unpaired, in case 4, *p* is paired with *A_t_(r)*. Further, in cases 2 and 4, the position *p* is considered to be *consistent* while in cases 1 and 3, *p* is considered to be *inconsistent*. An example of an intermediate structure is shown in Table S1 and Figure 1B.
III. The intermediate structure is decomposed into basic structure elements: individual hairpins and stems (Figure 1C). Hairpins are identified first, then stems. Hairpins are identified by the following procedure:

1. the loops of the hairpins are identified first as base pairs with only unpaired nucleotides in between the pairing nucleotides.
2. From this base pair, both ends of the hairpin stem are extended until first base pair of a different hairpin is encountered on either end. The strands of the hairpin must contain the same number of pairing nucleotides. All single-strand nucleotides between pairing nucleotides are added to the hairpin as well.
3. If there are single-strand nucleotides following the last base pair of a hairpin, they are added to the hairpin while ensuring they are not shared by multiple neighboring hairpins. Stems are identified in between hairpins. Stems have two strands, the 5’ strand and the 3’ strand, identified by the following procedure:

1. the strands of the stem start at the first nucleotide not occupied by hairpins or previously identified stems at 5’ and 3’ ends of the intermediate structure for the 5’ and 3’ strands, respectively.
2. The strands are extended in opposite directions, i.e. the 5’ strand in 5’->3’ and the 3’ strand in 3’->5’ for the same number of base pairs, until a base pair belonging already to a hairpin is encountered. Unlike hairpins, stems are not extended with neighboring single-strand nucleotides.
IV. Identification of inconsistency of the elementary structure elements (Figure 1D). Structural elements are considered inconsistent, if their proportion of inconsistent positions identified in step II is over a given threshold. The threshold was set to 20% for hairpins and 10% for stems. The threshold values were identified based on optimization using both the cross-validation and large scale datasets.
V. De novo prediction of the structure of the inconsistent elements (Figure 1E). RNAfold and RNAduplex (8) were used for hairpins and stems, respectively. The goal of this step is that the prediction corrects the wrong structure information at inconsistent positions. The advantage is that the elements are small and therefore the prediction of their structure is highly reliable in contrast to the prediction of the whole structure.
VI. The de novo predicted structures of the inconsistent elements are combined (pasted) with the intermediate structure of the consistent elements (Figure 1F) to form the resulting structure.

**Figure 1.**
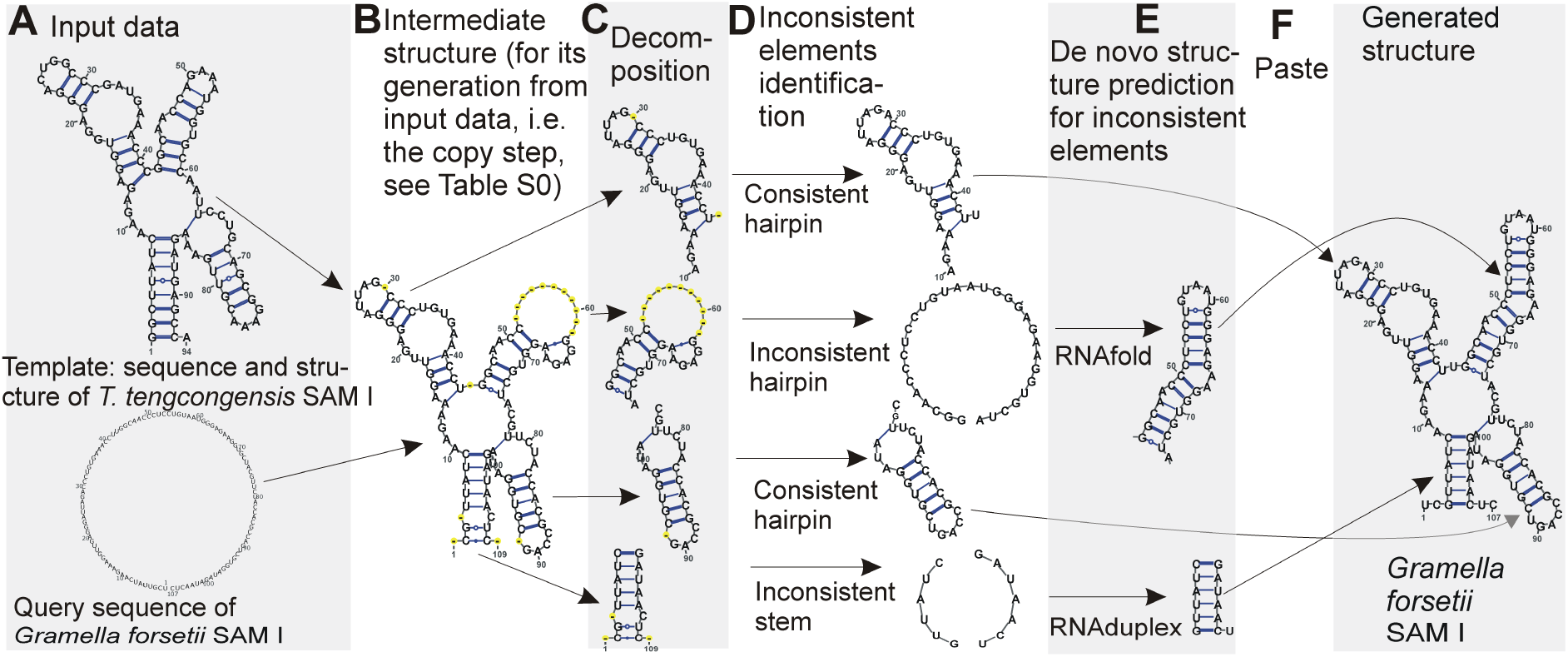
Demonstration of the method using SAM I structure. Structures are plotted by VARNA viewer (9). The sequence representation of the copy step is in Table S1. Note that VARNA interprets the signs for false non-canonical base pairs (‘3’), gapped base pairs (‘1’) and gaps (‘-‘) used in Table S1 as ‘-‘ signs in yellow circles.

### Bootstrap of the generated structure

Since the presented method will generate a structure for any input, even if the template and query sequences are completely unrelated, it is important that we distinguish reliable results from spurious ones.

We compute the reliability using a bootstrapping scheme. We use the query sequence to generate *N* sequences with randomly shuffled dinucleotides. For the shuffled sequences, structures are generated with the same procedure as for the query sequence.

First we validated a criterion that is evaluated by the bootstrapping scheme. We chose between tree edit distances and free energy (FE). For the first, distances *d_rnd_* = {*d*_*rnd*,*1*_,…,*d*_*rnd*,*N*_} between the generated structures and the template structure are computed. For the later, FEs *e*_*rnd*_ = {*e*_*rnd*,*1*_,…,*e*_*rnd*,*N*_} of the generated structures are computed.

Now *d_rnd_* and *e*_*rnd*_ approximate the distributions of tree edit distances and FEs obtained from non-homologous, i.e. shuffled sequences with the same length and nucleotide composition. The quality of the generated structure is then assessed with a z-score (10) relative to the population of non-homologous sequences:

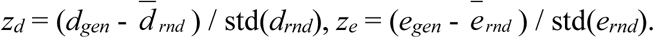

The generated structure of the original sequence is considered reliable with a *z*-score ≥ 2 (corresponding to the limit of the statistical significance of *p*-value = 0.05). In our experiments, we used *N* = 100.

Unlike direct use of the tree edit distance, the z-scores are relevant also when the query sequence is only a fragment of the template sequence. The generated structure is then naturally dissimilar to the template structure and has a relatively large tree edit distance. But reliable structure can still be generated by transferring the relevant substructure of the template. The presented bootstrapping scheme correctly classifies such substructures as reliable.

For purpose of validation of the bootstrap metrics and evaluation of the variance of z-scores, we repeated the bootstrap 100 times (100 runs with 100 randomized sequences each) for the 52 generated structures of the cross-validation dataset.

### Comparison to available methods

The presented method was compared to both a classical prediction method (RNAfold) and constrained prediction with RNAfold and refold.pl. The Vienna RNA package ver. 2.3.3 was used. The first method represents state-of-the-art of *de novo* prediction and is included mainly to put the improvements made by our method into proper scale. The latter tool should in theory perform the same task as our method and uses the same input information and thus is a more fair comparison.

The refold.pl script takes as input an alignment and a consensus structure. To perform template-based prediction we pass it a pairwise alignment between the subject sequence and the sequence of the template and the template structure extended to have the same length as the alignment by introducing the gaps identified by the alignment into it. The constraints were then used with `RNAfold –C`, as described in the Vienna RNA package user guide.

The comparison consists of two steps: first we validate the proposed method on a small dataset of experimentally identified RNA structures and then we perform a large scale evaluation on sequences without known structure.

### Cross-validation using experimentally identified structures

For the cross-validation, RNA families with at least two homologs with experimentally identified structures were identified and used. They let us to generate structure of one homolog using structure of other homolog of the same family as a template/constraint and vice versa. The generated structures were then compared to their experimentally identified counterparts.

The sequences and structures of the experimentally identified RNAs used for the cross-validation are shown in supplementary file S1.fasta. We collected 34 families with at least two experimentally identified structures per one family mainly from PDB, allowing for 52 predictions. The sources of the structures including databases and/or related papers are included in Table S4.

Accuracy of the generated/predicted structures was evaluated using two criteria: 1) percent of nucleotide positions with correctly predicted structural information, 2) tree edit distances (computed by RNAdistance (8)) to the experimentally identified structures. Ideally, the generated/predicted structures should have 100% of nucleotide positions with correctly predicted structural information and their tree edit distance to the experimentally identified structures should be zero.

### Large scale evaluation

The comparison was carried out using a reference dataset. Its characteristics are summarized in Table S2 and S3. The sequences are included in supplementary file S2.fasta. Templates including their sequences and structures are included in supplementary file S2a.fasta. The dataset was created from the test dataset of CentroidHomfold (11, 12) and extended with other RNAs to get more sequence/structure variability. The dataset consisted of 32 RNA families where at least one experimentally identified structure is known. In total, the dataset contains 3192 sequences with pairwise sequence similarity within families ranging from 43 to 95% and sequence similarity to the templates of 38 – 93%. The sequences were downloaded from Rfam (13) and SRPDB (14) databases, or when unavailable in the databases, identified using corresponding papers cited in Table S3 and downloaded from Genbank. As templates, the experimentally identified structures were used, downloaded together with their sequences from databases (mostly PDB (15)) or acquired using corresponding papers (Table S3).

## RESULTS

### Cross-validation with experimentally identified structures

The results of the cross-validation are summarized in Table S4 and Figure 2. The methodology of the cross-validation is explained in details in Methods. The proposed method generated more accurate structures than RNAfold and the refold method for 49 of total 52 predictions. The result was the same, when the accuracy was evaluated by tree edit distance and percentage of nucleotide positions with correctly predicted structural information.

**Figure 2.**
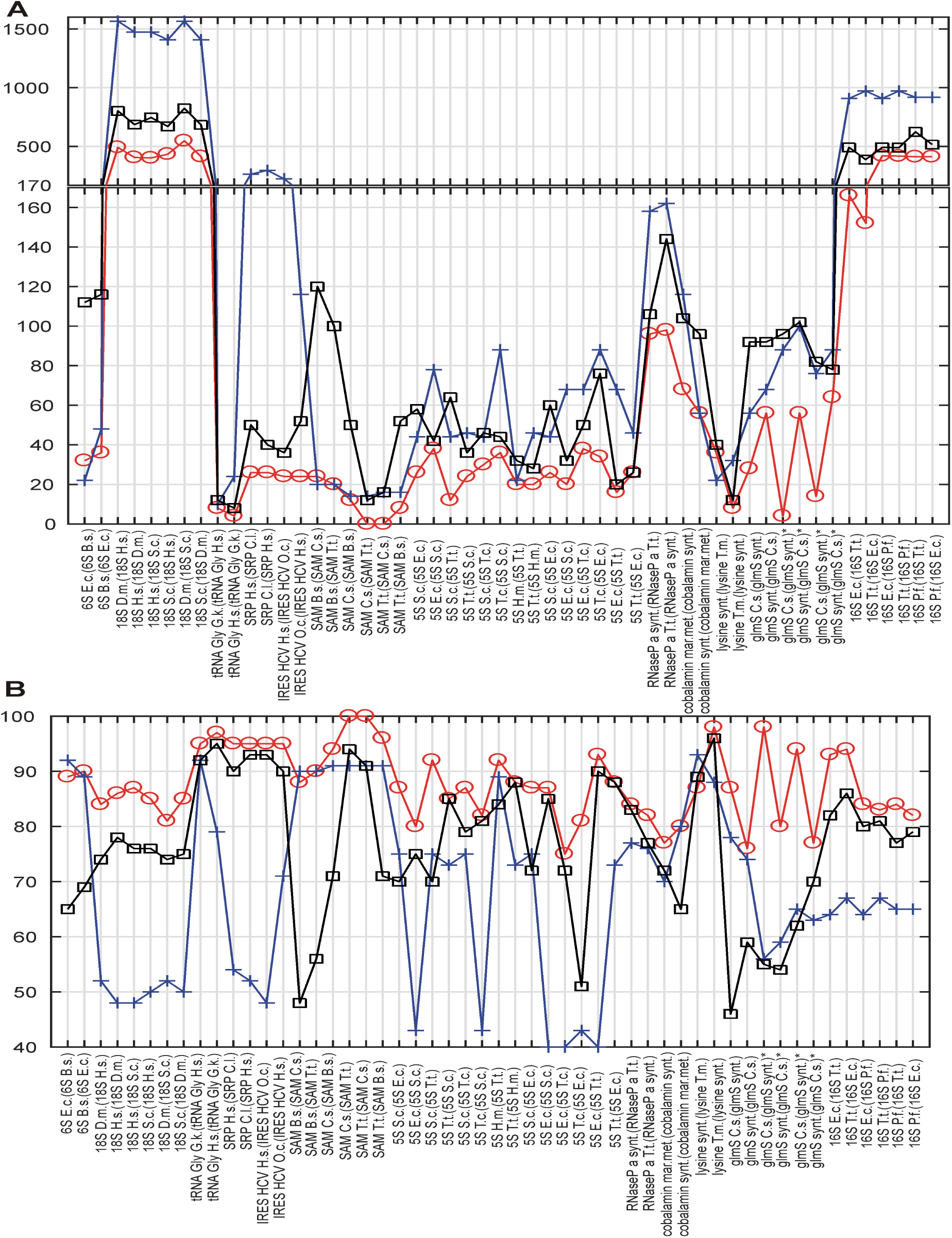
Cross-validation. In A and B, x-axis shows RNAs, whose structure is predicted and (in parenthesis) the RNAs, whose structures were used either for constraints or as templates. Y-axis shows tree edit distances (A) and percentage of nucleotide positions with correctly predicted structural information (B). Circles, squares and crosses show values for the proposed method, the refold method and RNAfold, respectively. For (A), the lesser the distance, the higher the structural similarity to experimentally identified structure; 0 for identical structures. For (B), the maximum of structural similarity to experimentally identified structure is indicated by 100%. For predictions marked with ∗, three structural versions were obtained by removing pseudoknots. Organism names are abbreviated: E.c.-*E*. *coli*, H.m.-*H*. *marismortui*, B.s.-*B*. *subtilis*, D.m.-*D*. *melanogaster*, H.s.-*H*. *sapiens*, O.c.-O. cuniculus, T.t.-*T*. *tencogensis*, C.s.-*C*. *subterraneus*, P.f.-*P*. *falciparum*, T.m.-*T*. *maritima*., S.c.-*S*. *coelicolor*.Figure 2 indicates that the proposed method was capable to generate more accurate structures than both classical prediction represented by RNAfold and the principally same method (the refold method). The main improvement is the ability to generate both large structures of long sequences and structures with long single-strand segments that are notoriously hard to be predicted with available prediction methods. Such typical structures here are rRNAs (16S and 18S rRNAs) and bacterial RNaseP. For some species of shorter and highly paired structures, as *E*. *coli* and *T*. *tencogensis* 5S RNAs and lysine riboswitches, the proposed method and the refold method provided similar accuracy (Figure 2A and B). For three RNAs, namely *E*. *coli* 6S RNA, synthetic lysine riboswitch and *H*. *marismortui* 5S RNA, RNAfold was more accurate than both the proposed method and the refold method. These structures were both shorter (with less than 190 nucleotides) and highly paired, which made them convenient for RNAfold prediction.

With respect to the structure similarity metrics, both tree edit distance and percentage of nucleotide position with correctly predicted structural information, were similarly reliable, indicating that tree edit distance is a good indicator of structure quality. We use tree edit distance in the remainder of the evaluation as the other metrics depends on the method used for sequence alignment. Note that the value of the tree edit distance depends on the size of structures and, as it is a distance: the higher the similarity, the lower the score, and the value of zero indicates structural identity.

### Reliability of the generated structure

The bootstrap procedure for evaluation of the reliability of the generated structure and its metrics (see Methods) was validated using the cross-validation dataset, i.e. experimentally identified structures. The FE-based z-scores obtained by the repeated bootstrap (100 runs with 100 randomized sequences each) evaluated 18 of total 52 generated structures as unreliable. Nevertheless, 15 of these 18 unreliable structures were false negatives (FNs). An example of a FN is shown Figure 3. It is the secondary structure of *C*. *subterraneus* glmS ribozyme generated using synthetic glmS ribozyme as the template. The generated structure is obviously accurate (cf. Figure 3B and D), but its z_e_ = 1.85, marking it as unreliable (z_e_ < 2). For comparison, we generated true negative (TN) structure of the same sequence using the proposed method with different values of inconsistency thresholds (30% for hairpins and 20% for stems) (Figure 3C). Its z_e_ = 2.7, i.e. evaluated as reliable (> 2), though it is obviously unreliable (cf. Figure 3C and B) and therefore a false positive. The accuracy of both the structures was documented well by their tree edit distances (d=4 and d=44, respectively) to their experimentally identified counterpart. Analogous situations occurred for the other 15 generated structures that were evaluated as unreliable by the FE-based z-scores.

**Figure 3.**
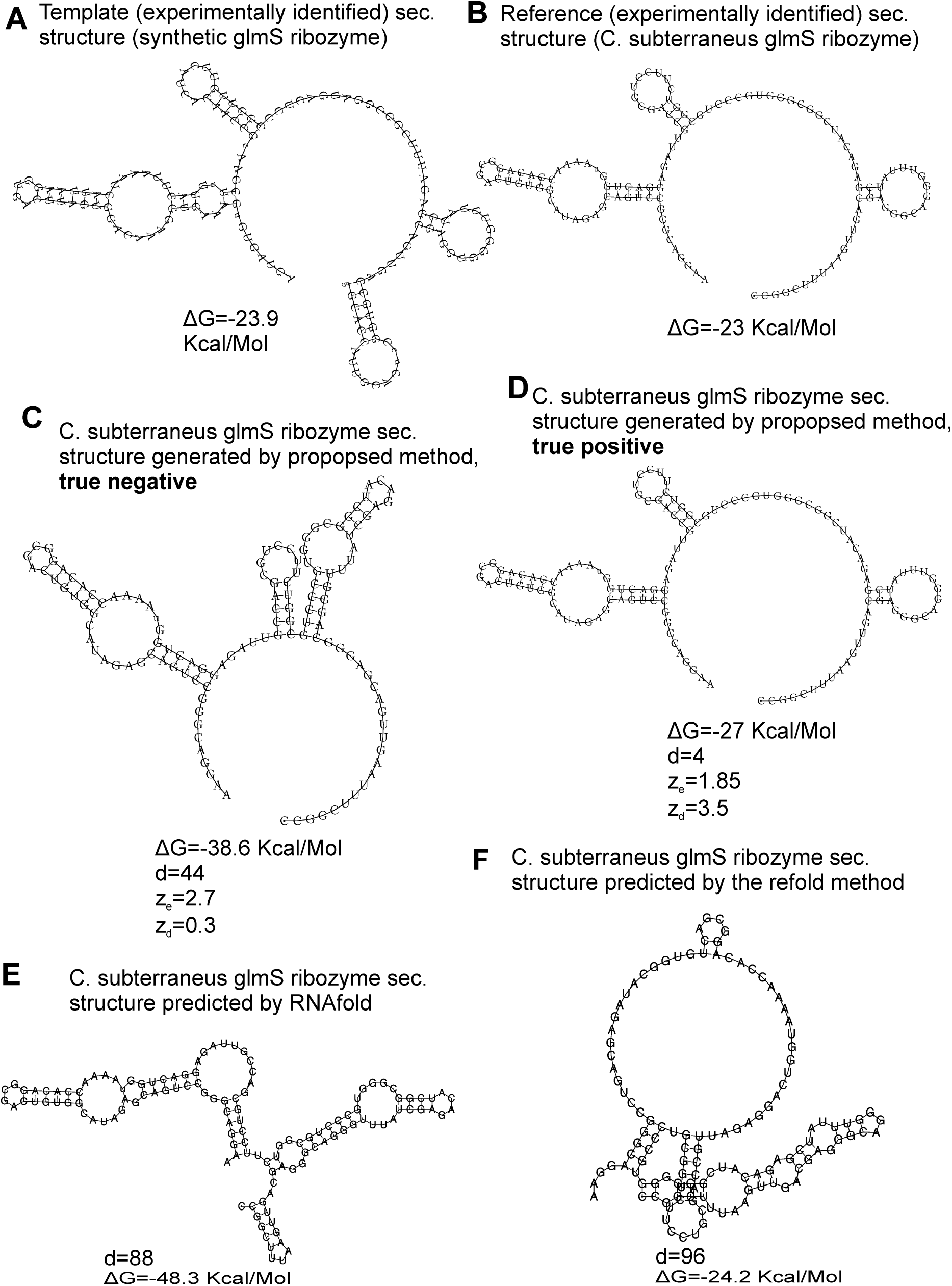
Validation of the structural similarity metrics. An example of structure of *C*. *subterraneus* glmS ribozyme is used for demonstration. Its experimentally identified secondary structure (derived from PDB ID 3b4c) is shown in A. The structure was predicted by RNAfold (B), the refold method (C) and the proposed method (D). For the latter two, experimentally identified secondary structure of homologous synthetic glmS ribozyme (PDB ID 3l3c)(shown in E) was used for constraint and as a template, respectively. ΔG – free energy, d - tree edit distance, per - percentage of nucleotide positions with correctly predicted structural information.

We therefore used tree edit distance between generated structures and templates instead of FE. The distance-based z-scores evaluated 3 of 52 generated structures as unreliable with more than 50% of their z-scores obtained by the repeated bootstrap (100 runs with 100 randomized sequences each) less than 2.

Our z-scores are however not definitive proofs of quality of the structure and z > 2 should be interpreted only as a high likelihood of the structure is reasonable. The main reason is the z-scores’s variance with respect to the randomized structures used in the bootstrap. The variance was estimated using the repeated bootstrap (100 runs with 100 randomized sequences each) for the 52 generated structures of the cross-validation dataset and counting how many z-scores were < 2 and > 2 for each generated structure. Besides of the 3 unreliable structures, 39 of total 52 structures had 100% z-scores greater than 2. Remaining 10 structures had their z-scores greater than 2 from 90.6% in average (for individual values, see Figure S1, the black curve).

For the example in Figure 3, the z-scores were z_d_=3.5 and 0.3 for the accurate and inaccurate structures, respectively (Figure 3D and C, respectively). Such z-scores better corresponded to the reliability of the structures.

The reason why FE was inadequate for our task was most likely its position independency. Two dissimilar structures with similar base pairs, though at different position on a sequence, can have similar FEs. As a result, an inaccurate structure can have correct FE, as demonstrated by the example in Figure 3. This fact is further demonstrated by the structure predicted by the refold method (Figure 3F) that is relatively dissimilar to its experimentally identified counterpart (Figure 3B). Nevertheless, the difference in FE between the predicted and experimentally identified structure is 1.2 Kcal/Molecule (-23 Kcal/Mol - -24.2 Kcal/Mol). The structure generated by the proposed method (Figure 3D) is fairly similar to the experimentally identified structure, but the difference in FE is 3.8 Kcal/Mol (-23 Kcal/Mol - -27 Kcal/Mol), i.e. higher, indicating stronger dissimilarity than for the structure predicted by the refold method.

### Large scale evaluation

As shown above, tree edit distance is biologically more relevant for comparison than free energy. In the following section we thus treat tree edit distance as our primary metric.

Results of the large-scale evaluation are summarized by Table S5 and Figure 4. They both document higher accuracy of the presented method when compared to both RNAfold and the refold-based method. In the first case, the higher accuracy was achieved due to the extra information used by the presented method. For the latter, that uses the same input information, the higher accuracy was is due to the active search for inconsistent structural elements and correction of their structure.

**Figure 4.**
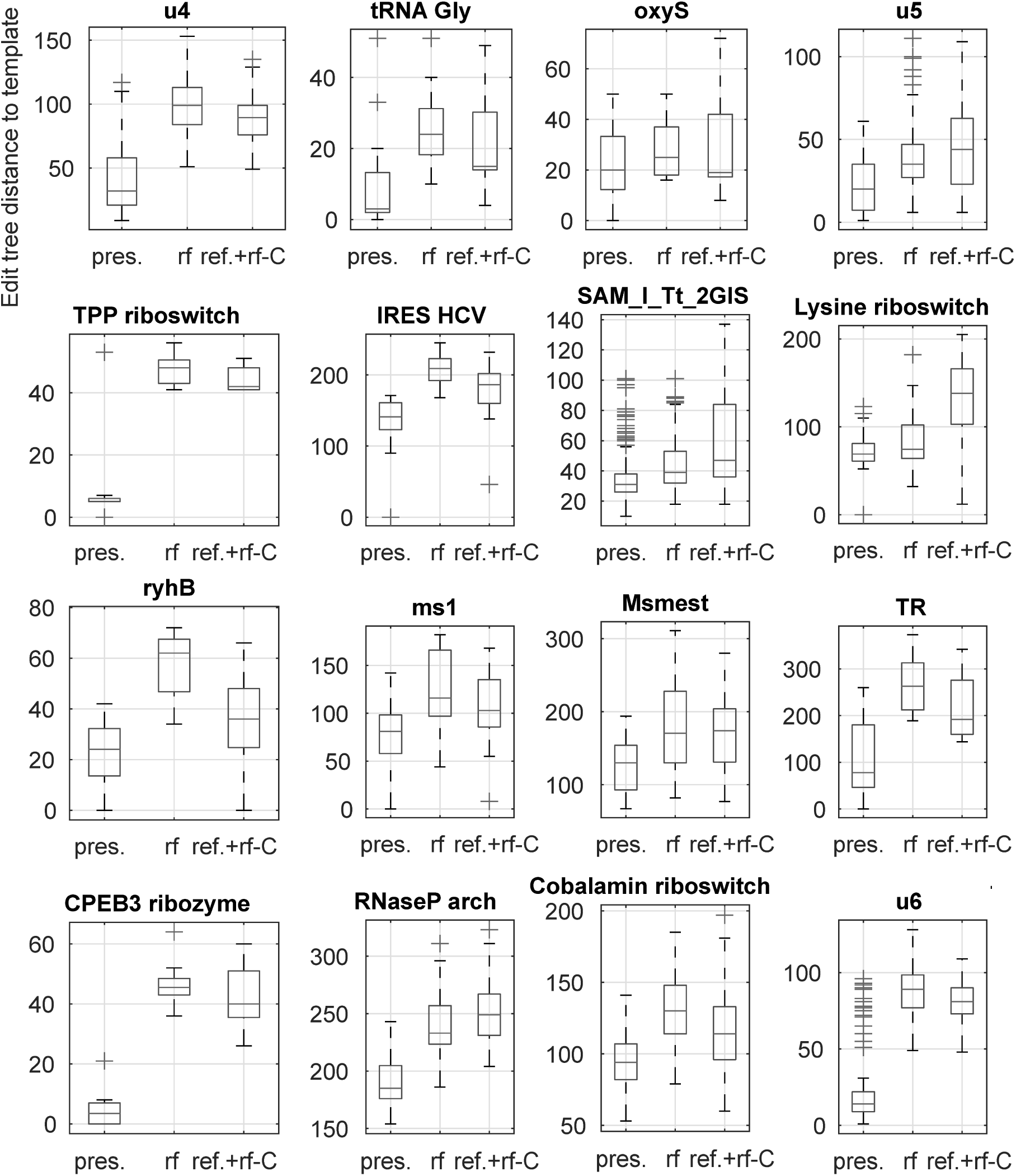
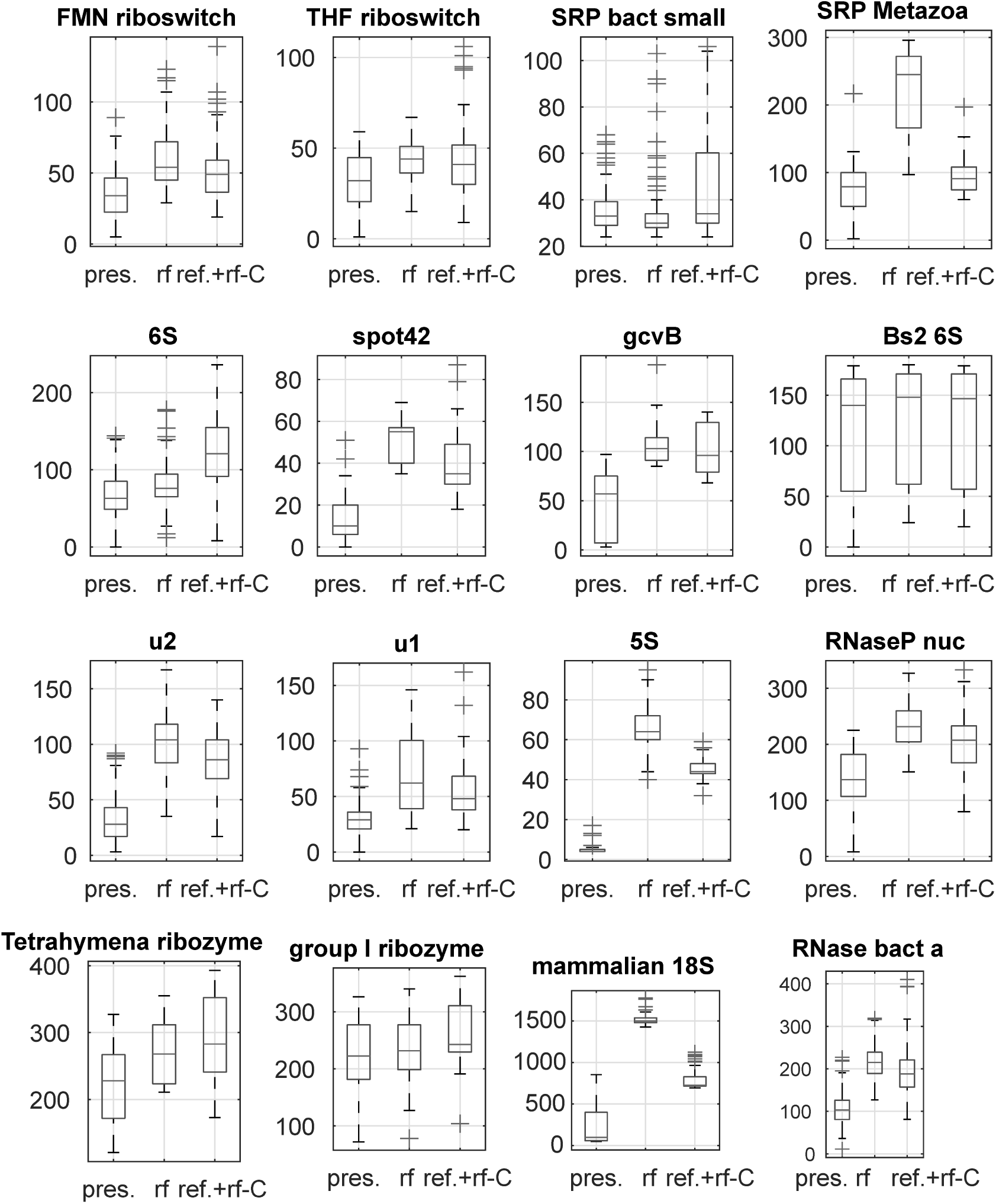
Comparison of the presented method. The compared methods were RNAfold as a representative of classical, single sequence secondary RNA structure prediction, and a refold.pl-based method that allows for the principally same type of prediction as the presented method. In the figure, 32 panels show results for 32 families of the reference dataset. In each panel, three box plots for the refold method, RNAfold and the presented method are shown (x-axis). Individual box plots show the median (red line), the 25th and 75th percentiles (the tops and bottoms of the boxes, respectively) and outliers (the whiskers) of the edit tree distances of the predicted structures of a single family to the templates. The distances between the tops and bottoms of the boxes are the interquartile ranges. The families are indicated by titles of the plots.

For 3 families (SRP bact small, Bs2 6S and group I ribozyme) RNAfold and the proposed method performed nearly the same. These families include densely and unambiguously paired structures that are convenient for the classical prediction, represented by RNAfold.

### Examples

In the following, the presented method is demonstrated in details using selected RNAs from the reference dataset. The examples are intended to illustrate situations when the proposed method is advantageous. Additional state-of-the-art prediction algorithms, principally different from the proposed method, were included in this demonstration to cover a broader spectrum of available prediction methods.

### Large single-strand segments: gcvB RNA

The first example is gcvB RNA, whose structure, experimentally identified in Sharma et al., 2007 (16) for *S*. *typhimurium* (Figure 5A), is difficult to predict as it includes relatively long single-strand segments. For this example we predict structures of gcvB homologs identified in the above cited paper. The sequence and structure of the template and the sequences of gcvB homologs and their predicted/generated structures are in supplementary file S3.fasta in sections a) and b), respectively.

**Figure 5.**
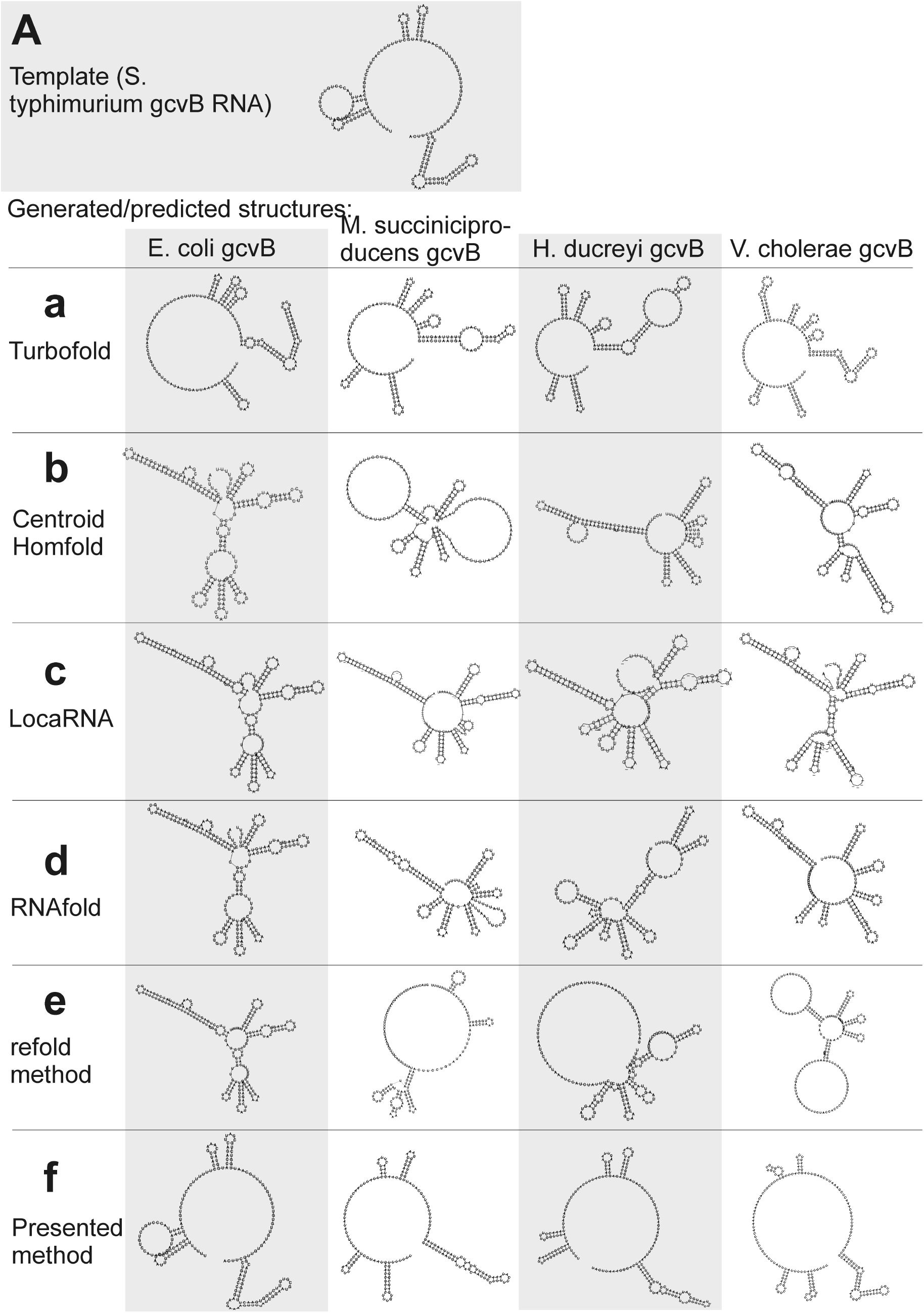
Individual secondary structures of gcvB RNA homologs predicted by available methods (a-e) and generated by the presented method (f). Structures are organized in rows and columns according to the method and species, respectively. The experimentally identified template structure of *S*. *typhimurium* gcvB RNA structure is shown at the top.

RNAfold and CentroidHomfold (used for single sequence prediction) tended to pair the sequences of long single-strand segments (Figure 5b and d). More accurate was Turbofold thanks to all the sequences of the homologs used as input (Figure 5a). Locarna and the refold.pl-based method that used the experimentally identified structure as constraint did not predict plausible structures of homologs (Figure 5c and e). Tree edit distances that quantify the similarity of the generated/predicted structures areshown in Table S6.

In contrast, the presented method was capable to generate structures that were similar to the experimentally identified structure (cf. Figure 5f and Figure 5A). The similarity is measured by tree edit distances (see Table S6). It was for two reasons: (i) the wrong, excessive pairing was prevented by the information of single-stranded segments copied from the template. This made the proposed method more accurate than the single-sequence prediction methods. (ii) The proposed method actively searched for inconsistent structural elements after copy step and predicted their structure *de novo*. This made it more accurate than the other methods that use the same information as input. Comparison of the accuracy using tree edit distances is in Table S6.

The improved accuracy can help to recognize non-homologous sequences. It is demonstrated here with the sequence of *E*. *coli* gcvB with randomly shuffled dinucleotides. Structure of this shuffled sequence, which represented an RNA not homologous to gcvB, could be distinguished from the gcvB homologs by its edit tree distance to the experimentally identified gcvB structure, when generated by the proposed method. The distance was twice longer than those of the structures of the gcvB homologs (Table S6). Recognition of this non-homologous RNA was not clear by the available methods as its tree edit distance was not unambiguously higher than the distances of the gcvB homologs (Table S6). The predicted structures of the shuffled RNA are in supplementary file S4.fasta.

The non-homologous RNA with the shuffled sequence could also be recognized by z-score of its generated structure. It was -3.6, which indicated strong unreliability. In contrast, z-scores for the gcvB homologs were all higher than 2 (namely 9.7, 3.6, 2.9 and 4.4 for *E*. *coli*, *V*. *cholera*, *H*. *ducreyi* and *M*. *succiniciproducens*, respectively). The usefulness of z-scores of the generated structures is further demonstrated in the next example.

### Large structure: 18S ribosomal RNAs

Large structures with long sequences are another class of sequences, when the classical prediction is often inaccurate. This is demonstrated here by the structure of mammalian 18S rRNAs. The methods of classical prediction that use either a single input sequence (RNAfold) or multiple input sequences (CentroidHomfold, Turbofold), and also methods that use a homologous experimentally identified structure (*H*. *sapiens* 18S rRNA) as a constraint (Locarna and the refold method) were largely inaccurate. This is demonstrated visually by Figure 6b, c (for RNAfold and the refold method only from technical reasons due to the large size of the 18S rRNA structures). The predicted structures were included in supplementary file S5.fasta.

**Figure 6.**
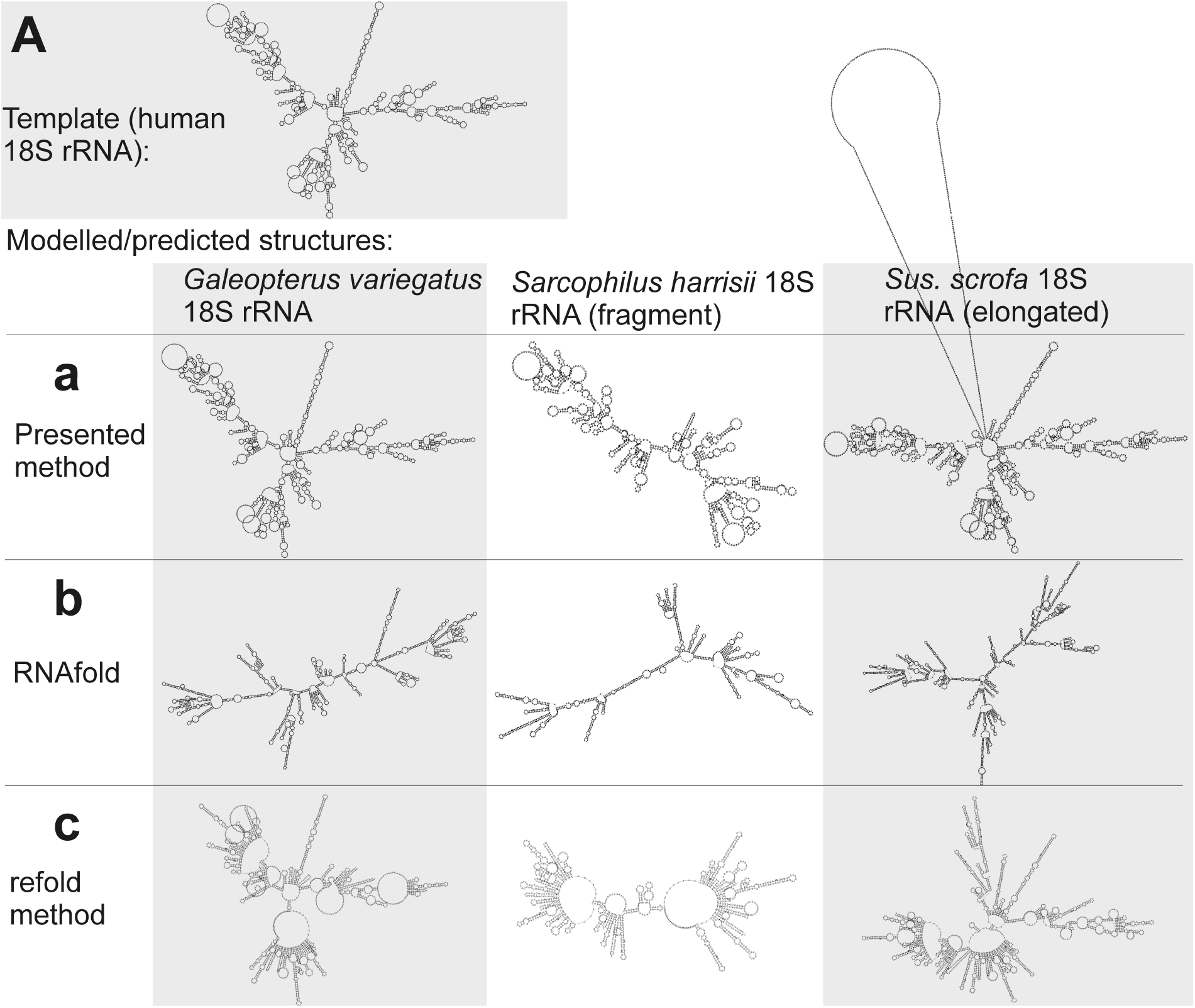
Individual secondary structures of 18S rRNA homologs predicted by available methods (b-c) and generated by the presented method (a). Structures are organized in rows and columns according to the method and species, respectively. The experimentally identified template structure of *H*. *sapiens 18S rRNA* structure is shown at the top.

The presented method was more accurate as its accuracy is largely independent of sequence length (Figure 6a). The improved accuracy was demonstrated by shorter tree edit distances of the generated structures to the experimental identified template of *H*. *sapiens* 18S rRNA (Table S7).

Interesting is the identification of the 18S rRNA fragment of *S*. *harrisii* 18S rRNA and the elongated 18S sequence of *S*. *scrofa* 18S rRNA. The proposed method identified correctly that the *S*. *harrisii* fragment contained only the expansion segments 3 and 6 of the whole 18S rRNA structure (Figure 6a). In contrast, the *S*. *scrofa* sequence included, beside the entire 18S rRNA structure, an additional ~700 nucleotides flanking the regular 18S rRNA structure (Figure 6a). Tree edit distances of these two structures were relatively high, when compared to the *G*. *variegatus* 18S rRNA structure that is complete (Table S7). This was due to the natural dissimilarity of either structural fragments or elongated structures to regular structures. However, z-scores were far greater than 2 (6.6, 12.1 and 19.2 for *S*. *harrisii*, *S*. *scrofa* and *G*. *variegatus*, respectively) indicating that these sequences are genuine 18S rRNAs, yet fragmented/elongated.

An interesting experiment and also validation of the above identification was to use an RNA that was non-homologous to both the fragment and the elongated sequence. To that end, we deployed *E*. *coli* 16S rRNA as a template. Now the template and the query sequences were not longer homologous, yet still related (all were rRNAs), and the z-scores should indicate unreliability of the generated structures. Indeed, the z-scores were -0.2, -6 and 0.2 for *S*. *harrisii*, *S*. *scrofa* and *G*. *variegatus*, respectively, indicating that the query sequences were not homologous to the 16S rRNA. In general, this procedure makes it possible to recognize, when the template and query sequences are not homologous, in other words, when the transfer of the template structure is biologically irrelevant producing wrong structures. What is important is that this bootstrap-based procedure is independent of the fact that query sequences are fragmented or elongated, as demonstrated in both this example and the previous example.

## DISCUSSION AND CONCLUSIONS

A method for template–based prediction/generation of single-sequence secondary RNA structure is presented. As demonstrated on examples, it is useful for determining whether an RNA molecule under investigation can conform to a secondary structure taken from a different molecule. This is useful for both obtaining RNA secondary structures and estimating ability of sequences to adopt the investigated structure. The method provides a solution in situations when available methods for secondary RNA structure prediction can not be used or are inaccurate.

It does not mean that a new *de novo* RNA secondary structure prediction algorithm/method is devised. It should be stressed that the presented method requires sequences and structures as input in contrast to the prediction methods that usually work only with sequences. However, when experimentally derived structures of homologous sequences are available, our method is able to correctly predict true biological structures, as shown by our cross validation study. We have also performed a large scale comparison of the results provided by our method for sequences where the ground truth is unknown, where our method also performed favorably. The large scale comparison is not used to benchmark the method, but merely for establishing a basis for evaluating the efficiency of the presented method.

Our method fills a gap resulting from the poor performance of the available methods of constrained prediction using known structures. Such prediction is increasingly useful as number of experimentally identified RNA structures (and thus the number of available templates) grows. The presented method is useful in various situations as demonstrated in this work. The method does not depend on length of sequences neither on the type of structure. It is fast and robust and it can be used for characterization of large numbers of sequences including fragments by structures of other RNAs. It produces z-scores based on bootstrapping of generated secondary structures that indicate whether the generated structures are relevant for the sequences. A Matlab-compiled executable is available on request.

## REFERENCES

D. H. Mathews: Revolutions in RNA secondary structure prediction. J Mol Biol, 359(3), 526-32 (2006) doi:S0022-2836(06)00108-2 [pii] 10.1016/j.jmb.2006.01.067

P. P. Gardner and R. Giegerich: A comprehensive comparison of comparative RNA structure prediction approaches. BMC Bioinformatics, 5, 140 (2004) doi:10.1186/1471-2105-5-140 1471-2105-5-140 [pii]

R. Lorenz, S. H. Bernhart, C. Honer Zu Siederdissen, H. Tafer, C. Flamm, P. F. Stadler and I. L. Hofacker: ViennaRNA Package 2.0. Algorithms Mol Biol, 6, 26 (2011) doi:1748-7188-6-26 [pii] 10.1186/1748-7188-6-26

D. H. Mathews: Using an RNA secondary structure partition function to determine confidence in base pairs predicted by free energy minimization. RNA, 10(8), 1178-90 (2004) doi:10.1261/rna.7650904 10/8/1178 [pii]

C. Smith, S. Heyne, A. S. Richter, S. Will and R. Backofen: Freiburg RNA Tools: a web server integrating INTARNA, EXPARNA and LOCARNA. Nucleic Acids Res, 38(Web Server issue), W373-7 (2010) doi:gkq316 [pii] 10.1093/nar/gkq316

R. R. Gutell, J. C. Lee and J. J. Cannone: The accuracy of ribosomal RNA comparative structure models. Curr Opin Struct Biol, 12(3), 301-10 (2002) doi:S0959440X02003391 [pii]

D. Higgins, J. Thompson, T. Gibson, J. D. Thompson, D. G. Higgins and T. J. Gibson: CLUSTAL W: improving the sensitivity of progressive multiple sequence alignment through sequence weighting,position-specific gap penalties and weight matrix choice. Nucleic Acids Res., 22, 4673-4680 (1994)

I. L. Hofacker: RNA secondary structure analysis using the Vienna RNA package. Curr Protoc Bioinformatics, Chapter 12, Unit 12 2 (2004) doi:10.1002/0471250953.bi1202s04

K. Darty, A. Denise and Y. Ponty: VARNA: Interactive drawing and editing of the RNA secondary structure. Bioinformatics, 25(15), 1974-5 (2009) doi:btp250 [pii]10.1093/bioinformatics/btp250

E. Kreyszig: Advanced Engineering Mathematics. John Wiley & Sons Inc, (1979)

M. Hamada, K. Sato, H. Kiryu, T. Mituyama and K. Asai: Predictions of RNA secondary structure by combining homologous sequence information. Bioinformatics, 25(12), i330-8 (2009) doi:btp228 [pii] 10.1093/bioinformatics/btp228

H. Kiryu, T. Kin and K. Asai: Robust prediction of consensus secondary structures using averaged base pairing probability matrices. Bioinformatics, 23(4), 434-41 (2007) doi:btl636 [pii] 10.1093/bioinformatics/btl636

P. P. Gardner, J. Daub, J. G. Tate, E. P. Nawrocki, D. L. Kolbe, S. Lindgreen, A. C. Wilkinson, R. D. Finn, S. Griffiths-Jones, S. R. Eddy and A. Bateman: Rfam: updates to the RNA families database. Nucleic Acids Res, 37(Database issue), D136-40 (2009) doi:gkn766 [pii]10.1093/nar/gkn766

E. S. Andersen, M. A. Rosenblad, N. Larsen, J. C. Westergaard, J. Burks, I. K. Wower, J. Wower, J. Gorodkin, T. Samuelsson and C. Zwieb: The tmRDB and SRPDB resources. Nucleic Acids Res, 34(Database issue), D163-8 (2006) doi:34/suppl_1/D163 [pii]10.1093/nar/gkj142

J. L. Sussman, D. W. Lin, J. S. Jiang, N. O. Manning, J. Prilusky, O. Ritter and E. E. Abola: Protein Data Bank (PDB): Database of three-dimensional structural information of biological macromolecules. Acta Crystallographica Section D-Biological Crystallography, 54, 1078-1084 (1998) doi:Doi 10.1107/S0907444998009378

C. M. Sharma, F. Darfeuille, T. H. Plantinga and J. Vogel: A small RNA regulates multiple ABC transporter mRNAs by targeting C/A-rich elements inside and upstream of ribosome-binding sites. Genes Dev, 21(21), 2804-17 (2007) doi:21/21/2804 [pii]10.1101/gad.447207

